# Evolutionary divergence of basal and activity-dependent exon splicing in cortical neurons

**DOI:** 10.1101/2022.12.29.522197

**Authors:** Owen Dando, Jing Qiu, Siddharthan Chandran, Giles E. Hardingham

## Abstract

Alternative splicing of mRNA exons in mammalian neurons increases diversity of the proteome and is regulatable by signaling pathways. However, the degree of conservation of basal and signal-dependent exon usage between human neurons and those from experimental models such as mice is incompletely understood. We previously showed that cortical neuronal activity-dependent gene transcription exhibits human/mouse differences, driven by evolutionary divergence of cis-acting promoter elements (Qiu et al. 2016). Since alternative exon usage influences brain development and cognition, is controlled by neuronal activity, and is disturbed in brain disorders, we investigated human/mouse differences in exon usage in cortical neurons. Comparing orthologous exons, basal exon inclusion levels showed human-mouse conservation, but also significant differences determined by cis-acting sequences: human-mouse conservation and divergence in exon usage was recapitulated in neurons from Tc1 mice carrying human chromosome-21 (hCh21). Activity-dependent changes in exon usage also exhibited significant conservation: gene structure was more likely to be conserved in activity-regulated exons, and exons regulated in both human and mouse neurons were enriched in RBFOX and SAM68 targets, and genes were centred on cytoskeletal organisation, mRNA transcription/processing, and synaptic signaling. However, divergence was also evident, and human-specific activity-dependent exon usage was dominated by genes involved in lipid biosynthesis, signaling and trafficking. Notably, the pattern of activity-dependent usage of hCh21 exons in human neurons was not recapitulated in mouse Tc1 neurons. Thus, unlike species-specific differences in activity-dependent gene transcription, cis-acting DNA sequence divergence is insufficient to explain inter-species differences in activity-regulated exon usage. Trans-acting factors involved in activity-responsive splicing have likely also diverged.

## Introduction

Humans and mice diverged from their common ancestor approximately 80 million years ago. Nevertheless, over 90% of human genes have 1:1 orthologs in mice (Monaco et al., 2015). However, whether the expression of these orthologs in response to signaling pathways is conserved or otherwise was not well understood. A classic example of this type of response is found in CNS neurons, where electrical activity dynamically controls gene expression to influence neuronal development, neuroprotection, neurophysiological properties, and ultimately cognitive function. We previously reported that activity-responsiveness of expression levels of 1:1 orthologs in mouse and human cortical neurons showed evidence of evolutionary divergence (Qiu et al., 2016). The human system employed was glutamatergic cortical-patterned neurons from human embryonic stem cells (hESC^CORT^-neurons), and comparisons were made to mouse primary cortical neurons (Mus-PRIM^CORT^ neurons) at day in vitro (DIV) 4 and DIV10, as well as cortical-patterned neurons from mouse embryonic stem cells (Mus-ESC^CORT^-neurons). The rationale for employing multiple mouse neuronal preparations was to distinguish genuine evolutionary divergence from non-species-dependent differences such as developmental stage or cellular origin (primary tissue vs. stem cell). Mechanistically, we concluded that this divergence involved changes in cis-acting gene promoter regions that contain binding sites for activity-responsive transcription factors (Qiu et al., 2016). Other studies published shortly after also provided evidence of divergence of gene activity-responsiveness and showed that it could influence lineage-specific aspects of neuronal development (Ataman et al., 2016; Pruunsild et al., 2017), reviewed in Hardingham et al. (2018): (Hardingham et al., 2018).

While the basic property of whether a gene’s transcription is up-or down-regulated in neurons in response to electrical activity is of importance in determining the physiological outcome, so is post-transcriptional regulation. Eukaryotic genes possess coding exons interspersed with non-coding introns, the former of which are spliced to create protein-coding open reading frames. Many genes exhibit variable usage of exons (a form of alternative splicing) which enable a variety of related proteins to be encoded at a single genomic locus. Moreover, many exons are subject to signal-dependent inclusion (or exclusion). Neuronal electrical activity is known to control exon usage, and activity-dependent alternative splicing regulates neurophysiological and other properties by determining the function of specific proteins (Furlanis and Scheiffele, 2018). Aberrant alternative splicing in neurons, is thought to contribute to the pathogenesis of human brain disorders including Parkinson’s and Alzheimer’s diseases, and ALS-FTD (Li et al., 2021; Lopez Soto et al., 2019) and misregulation of activity-dependent exon usage is implicated in autism spectrum disorder phenotypes (Parikshak et al., 2016; Quesnel-Vallieres et al., 2016; Quesnel-Vallieres et al., 2019). However, despite mice being widely used to model human disease, the degree of conservation vs. divergence of alternative splicing in neurons from humans to mice is incompletely understood. Basal, steady-state alternative splicing within orthologs has diverged during evolution in many tissues and cell types (Verta and Jacobs, 2022), although studies on divergence in neurons, and of signal-dependent alternative splicing, are less advanced.

Here we have assessed the human-mouse divergence and conservation of orthologous exon splicing in cortical neurons, considering both ‘basal’ levels of exon inclusion as well as changes that occur in response to neuronal activity. We also wanted to identify the basis of any divergence: differences in cis-acting DNA sequences vs. trans-acting factors in the cellular environment (e.g. signaling molecules or splicing machinery). The Tc1 transchromosomic mouse strain carries a copy of human chromosome 21 (Hsa21), albeit with approximately 10% deleted (Gribble et al., 2013; O’Doherty et al., 2005). This enables basal and activity-dependent alternative splicing of human genes to be studied in a mouse neuronal environment, employing an *in silico* species-specific RNA-seq read sorting workflow we developed (Qiu et al., 2018). In both basal and activity-dependent alternative splicing we see significant conservation between mice and humans, but significant divergence too. Moreover, basal and activity-dependent alternative splicing of neuronal mRNAs have quite different requirements when it comes to the need for trans-acting factors to come from the correct species.

## Results and Discussion

### Evolutionary conservation vs. divergence of basal exon inclusion in cortical neurons

We first assessed evolutionary divergence in basal exon inclusion in human vs. mouse cortical neurons, before analysing any activity-dependent changes. We made genome-wide comparison between basal exon inclusion levels between Hum-ESC^CORT^ and DIV4 Mus-PRIM^CORT^ neurons (Qiu et al., 2016). Comparisons were restricted to “orthologous” exon inclusion/exclusion events, namely those whose upstream exon end, downstream exon start, and start and end of the alternatively spliced exon could be matched to within ten base-pairs, after translating co-ordinates between the mm10 and hg38 genome assemblies. We further restricted events to those whose average count, over all basal samples, of RNA-seq reads assigned to either inclusion or exclusion of the skipped exon, was greater than 5. We observed a significant correlation between Hum-ESC^CORT^ and DIV4 Mus-PRIM^CORT^ neurons when comparing the fractional exon inclusion (FEI) of orthologous exons (Fig. 1a,e). For each sample set we also classified every exon as primarily included (PI, FEI>0.8), primarily skipped (PS, FEI<0.2) or alternatively-spliced (AS, 0.8>FEI>0.2). In Hum-ESC^CORT^ neurons, alternatively-spliced exons were enriched 13-fold in those exons also alternatively-spliced in DIV4 Mus-PRIM^CORT^ neurons (Fig. S1a). Collectively this shows that the basal level of orthologous exon inclusion in human and mouse cortical neuronal mRNA transcripts exhibits significant evolutionary conservation, although the correlation was far from perfect. Fig. 1a (right) illustrates two orthologous exons (from *ZMYND11* and *HNRNPAB*), one of which (from *ZMYND1*) has a similar FEI in Hum-ESC^CORT^ and DIV4 Mus-PRIM^CORT^ neurons, and one (from *HNRNPAB*) is quite different.

**Figure 1.**
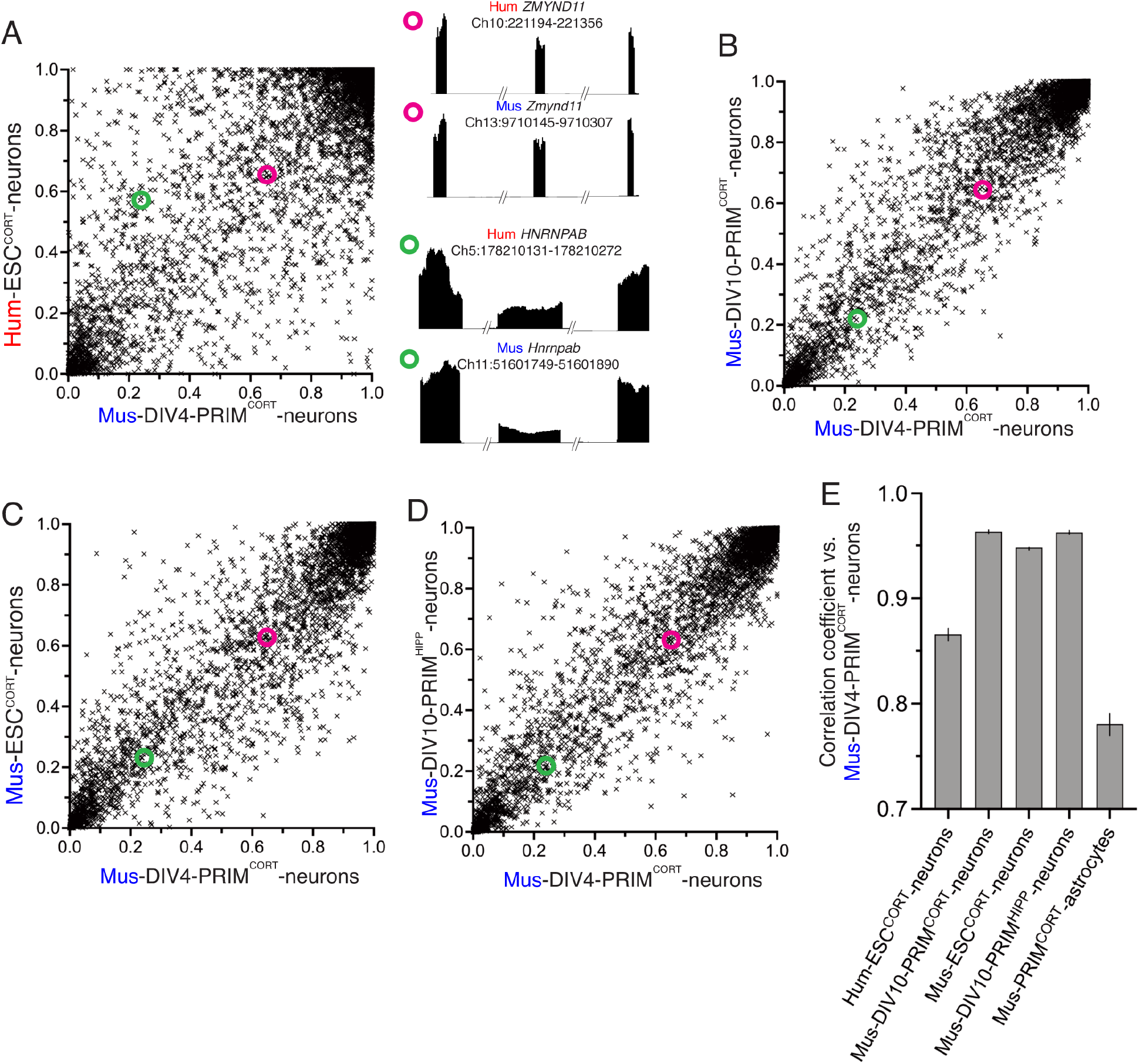
Evolutionary conservation vs. divergence of basal exon inclusion in forebrain neurons. **A-D)** Mean fractional exon inclusion (FEI) in DIV4 Mus-PRIM^CORT^ neurons plotted against the corresponding FEI in the indicated cell types (mean FEI, n=3 independent biological replicates here and throughout the figure). All exons plotted have a 1:1 human-mouse ortholog, the mean of 3 independent biological replicates is shown. (A, right) shows examples of two 1:1 orthologous exons (coordinates relate to this exon), plus flanking exons, showing relative RNA-seq read density. One exon (from *ZMYND11*) has a similar FEI in human and mouse neurons, while one (from *HNRNPAB*) has a different FEI in human and mouse neurons. The FEI of the *ZMYND11* exon and *HNRNPAB* exon is highlighted in the scatter graphs A-D with magenta and green circles, respectively. **E)** Pearson r correlation coefficients for the comparisons made in A-D, and Fig. S2b. Error bars indicate the 95% confidence limits and in all cases p<0.0001. For data relating to this figure see Source_Data.xlsx.

We wanted to determine whether evolutionary divergence in exon inclusion levels contributed to the differences. While this may be at first glance self-evident, other factors may be responsible, such as the two populations of neurons being at a different developmental stage, or derived from different sources (embryonic stem cell line vs. primary tissue), or even simple experimental variation. We therefore performed identical analyses between DIV4 Mus-PRIM^CORT^ neurons and more mature DIV10 Mus-PRIM^CORT^ neurons as well as with mouse ES cell-derived cortical neurons (Mus-ESC^CORT^-neurons) to determine the approximate influence of developmental stage (DIV4 vs DIV10) and tissue origin (primary vs. stem cell) on FEI. These inter-mouse comparisons showed a higher correlation between each other (Fig 1b,c,e) and higher enrichment of alternatively spliced genes than that observed in the Hum-ESC^CORT^ vs. DIV4 Mus-PRIM^CORT^ comparison. We also tested inter-lab replicability by comparing the FEI in our DIV4 Mus-PRIM^CORT^ neurons to that in DIV10 hippocampal (DIV10 Mus-PRIM^HIPP^) neurons generated and analysed by a different laboratory (Quesnel-Vallieres et al., 2016). We observed a good correlation: similar to that observed between mouse neuronal preparationd made in our laboratory (Fig. 1d,e). This suggests that maturation state and origin (tissue vs. stem cell) are unlikely to account for all of the changes in splicing observed between Hum-ESC^CORT^ and DIV4 Mus-PRIM^CORT^ neurons (Fig. 1a). Collectively this supports a model whereby basal FEI of orthologous exons in mRNAs from human vs. mouse cortical neurons exhibit significant evolutionary divergence. Interestingly though, mouse cortical neuronal exon usage pattern was found to be more similar to human cortical neurons than mouse cortical astrocytes (Fig. 1e, Fig S1b). This points to there being a neuron-associated genome-wide exon usage signature that distinguishes neurons from other neural cell types and is sufficiently conserved that exon usage in neurons from different mammalian species show greater correlation than different neural cell types (neurons and astrocytes) from the same species.

### Evolutionary conservation vs. divergence of activity-dependent alternative splicing

We next studied changes in exon inclusion within neurons in response to to L-type Ca^2+^ channel activation, an important mediator of activity-dependent gene regulation (Bito et al., 1997; Deisseroth et al., 2003; Ma et al., 2014; Sheng et al., 1990; West and Greenberg, 2011; Wheeler et al., 2012). To do this, hESC^CORT^-neurons and DIV4 Mus-PRIM^CORT^ neurons were treated ± KCl-induced membrane depolarization in the presence of the L-type Ca^2+^ channel agonist FPL64176, plus NMDA receptor antagonist MK-801 to prevent any excitotoxicity (hereafter KCl) for 4 h. After 4 h RNA was harvested and RNA-seq performed, followed by analysis of KCl-induced changes in exon usage. In both DIV4 Mus-PRIM^CORT^ neurons (Fig. 2a) and hESC^CORT^-neurons (Fig 2b) splicing levels changed significantly in 800-900 orthologous exons, with KCl treatment causing an increase in exclusion of certain exons, and inclusion of others. Others have reported previously that in mouse hippocampal neurons, activity-induced exclusion/skipping of exons in response to neuronal activity is more prevalent than activity-induced increases in exon inclusion (Quesnel-Vallieres et al., 2016). This is also something we observe, not only with mouse neurons but human neurons too (Fig. 2a,b). Similar analyses were performed for DIV10 Mus-PRIM^CORT^ neurons ± KCl (Fig. 2c), Mus-ESC^CORT^-neurons ± KCl (Fig 2d) and on RNA-seq data obtained by another laboratory: a 3h KCl stimulation of DIV10 Mus-PRIM^HIPP^ neurons (Quesnel-Vallieres et al., 2016) (Fig. 2e).

**Figure 2.**
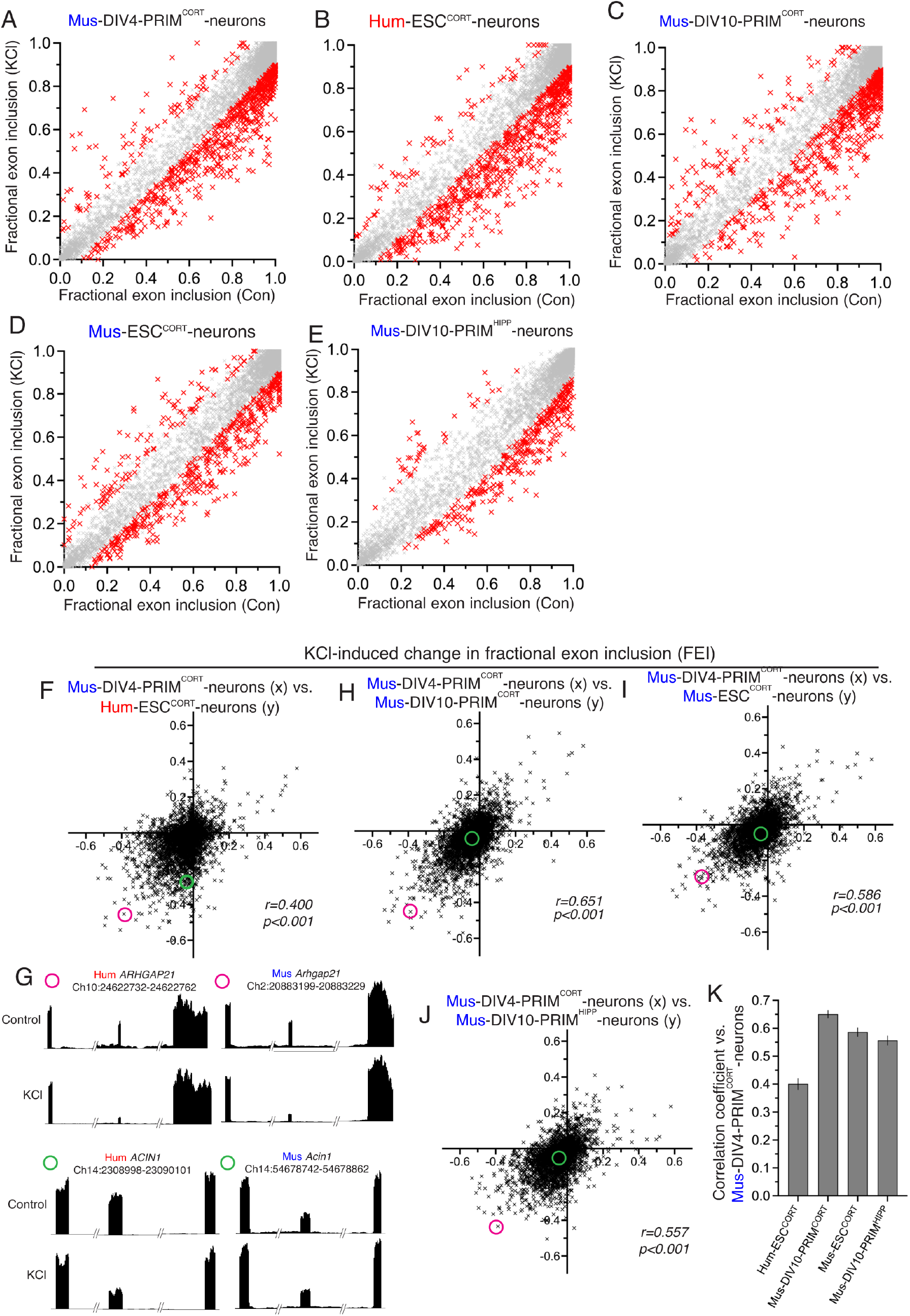
Evolutionary conservation vs. divergence of activity-dependent alternative splicing. **A-E)** For the indicated neuronal preparations, FEI in control neurons is plotted against KCl-stimulated neurons (A-D: 4h; E: 3h-data were generated by another lab (Quesnel-Vallieres et al., 2016)). All exons plotted have a 1:1 human-mouse ortholog. Red crosses indicate a significant difference in FEI (p<0.05, average read count for exon inclusion or exclusion over all samples > 5, inclusion level difference > 0.1, n=3). **F)** The KCl-induced change in FEI in DIV4 Mus-PRIM^CORT^ neurons is plotted against the corresponding change in Hum-ESC^CORT^ neurons. **G)** Examples of two 1:1 orthologous exons (coordinates relate to this exon), plus flanking exons, showing relative RNA-seq read density. One exon (from *ARHGAP21*) has a similar KCl-induced FEI in human and mouse neurons, while one exon (from *ACIN1*) is only subject to activity-dependent alternative splicing in human neurons. The KCl-induced FEI change of the *ARHGAP21* exon and *ACIN1* exon is highlighted in (F) with a magenta and a green circle, respectively. **H-J)** The KCl-induced change in FEI in DIV4 Mus-PRIM^CORT^ neurons is plotted against the corresponding change in the indicated cell types. The KCl-induced FEI change of the *ARHGAP21* exon and *ACIN1* exon described in 2G is highlighted with a magenta and a green circle, respectively. **K)** Correlation coefficients for the comparisons made in F-I. Error bars indicate the 95% confidence limits and in all cases p<0.0001 For data points relating to this figure see Source_Data.xlsx.

Plotting KCl-induced changes in FEI in hESC^CORT^ vs. DIV4 Mus-PRIM^CORT^ neurons revealed a correlation, albeit quite weak (Fig. 2f,k). Fig. 2g shows examples of RNA-seq read density in two 1:1 orthologous exons for both species ± KCl treatment. One exon (from *ARHGAP21*) has a similar KCl-induced FEI in human and mouse neurons, while one exon (from *ACIN1*) is only subject to activity-dependent alternative splicing in human neurons. Globally, the correlation between KCl-induced changes in FEI in DIV4 Mus-PRIM^CORT^ neurons vs. DIV10 Mus-PRIM^CORT^ neurons and vs. mESC^CORT^-neurons are significantly higher (Fig. 2h,i,k). We also compared KCl-induced changes in FEI DIV10 Mus-PRIM^CORT^ neurons with activity-dependent changes in FEI calculated from similar RNA-seq data obtained by another laboratory: a 3h KCl stimulation of DIV10 mouse hippocampal neurons (Quesnel-Vallieres et al., 2016), also revealing a correlation similar to our comparisons of DIV10 Mus-PRIM^CORT^ neurons vs. DIV4 Mus-PRIM^CORT^ neurons and mESC^CORT^-neurons (Fig. 2j,k). Thus, activity-dependent changes in FEI in mouse forebrain neurons show good inter-lab replicability and are relatively insensitive to developmental stage, tissue origin (primary vs stem cell) and also forebrain region (neocortex vs. hippocampus). Moreover there is evidence of human-mouse evolutionary divergence.

One factor to consider when assessing differences in KCl-induced changes in FEI in different neuronal preparations is that the magnitude of change may be influenced by differences in basal FEI, which we know shows human-mouse evolutionary differences (Fig. 1). For example, a mouse exon with a basal FEI of 0.4 that increases upon KCl stimulation has a theoretical maximal FEI change of 0.6, whereas if the orthologous human exon has a basal FEI of 0.7 then the maximal FEI change possible is only 0.3. We therefore performed additional comparisons of KCl-induced inclusion level differences calculated as a % of the maximum possible inclusion level difference. We restricted our analysis to orthologous exons where basal FEI was >0.2 and <0.8 to eliminate excessive skewing of the data caused by modest absolute changes in FEI giving very high percentage figures (e.g. an FEI changing from 0.95 to 0.98 would give a figure of 60%). Our comparisons (Fig. S2a-e) mirrored those made in Fig. 2f-j and yielded similar results: KCl-induced changes in exon inclusion in hESC^CORT^ vs. DIV4 Mus-PRIM^CORT^ neurons showed significant correlation, but it was weaker than when comparing KCl-induced changes in exon usage in DIV4 Mus-PRIM^CORT^ with any of the other mouse neuronal preparations (DIV10 Mus-PRIM^CORT^ neurons, mESC^CORT^-neurons, and DIV10 Mus-PRIM^HIPP^ neurons. These observations are consistent with their being evolutionary divergence in activity-dependent alternative splicing of orthologous exons between mice and humans.

### Ontology of human-specific vs. conserved activity-dependent alternatively spliced genes

There are several examples of activity-dependent changes in exon inclusion influencing the function of the protein encoded by the alternatively spliced transcript (Furlanis and Scheiffele, 2018). This can require that an exon encodes a functionally autonomous part of a protein so it can be included or excluded to alter a protein’s function without causing non-specific loss of function (e.g. through protein misfolding or removing part of a key structural domain). We reasoned that the organisation of exons in a gene is more likely to be conserved in those exons subject to signal-dependent regulation. Indeed, taking exons that are subject to activity-dependent regulation in mouse neurons (DIV4 and DIV10 Mus-PRIM^CORT^, mESC^CORT^-neurons) we observed that they are enriched in 1:1 human-mouse orthologs by 2.7-fold, indicating selective evolutionary pressure to maintain genetic structure.

Taking genes containing orthologous exons subject to activity-dependent changes in FEI in both human and mouse neurons, ontological analysis revealed three main functional areas (Fig. 3a-c). Prominent processes and functions are associated with cytoskeletal organisation and transport along cytoskeletal tracks (Fig. 3a). The second major area is in the control of gene expression including transcriptional control, epigenetic regulation, and post-transcriptional mRNA processing, including polyadenylation and RNA splicing itself (Fig. 3b). A third prominent area of conserved activity-responsive AS functions involve synaptic signaling and action potential firing (Fig. 3c), and associated subcellular components such as synaptic vesicles, the pre- and post-synaptic membrane, and specialist structures including the post-synaptic density, AMPA receptor complex and the axon initial segment. These processes have been highlighted recently as subject to regulation by activity-dependent AS, particularly in the context of homeostatic plasticity in mice (Iijima and Yoshimura, 2019; Thalhammer et al., 2020) and our data suggests that activity-dependent AS may play a similar role in human neurons.

**Figure 3.**
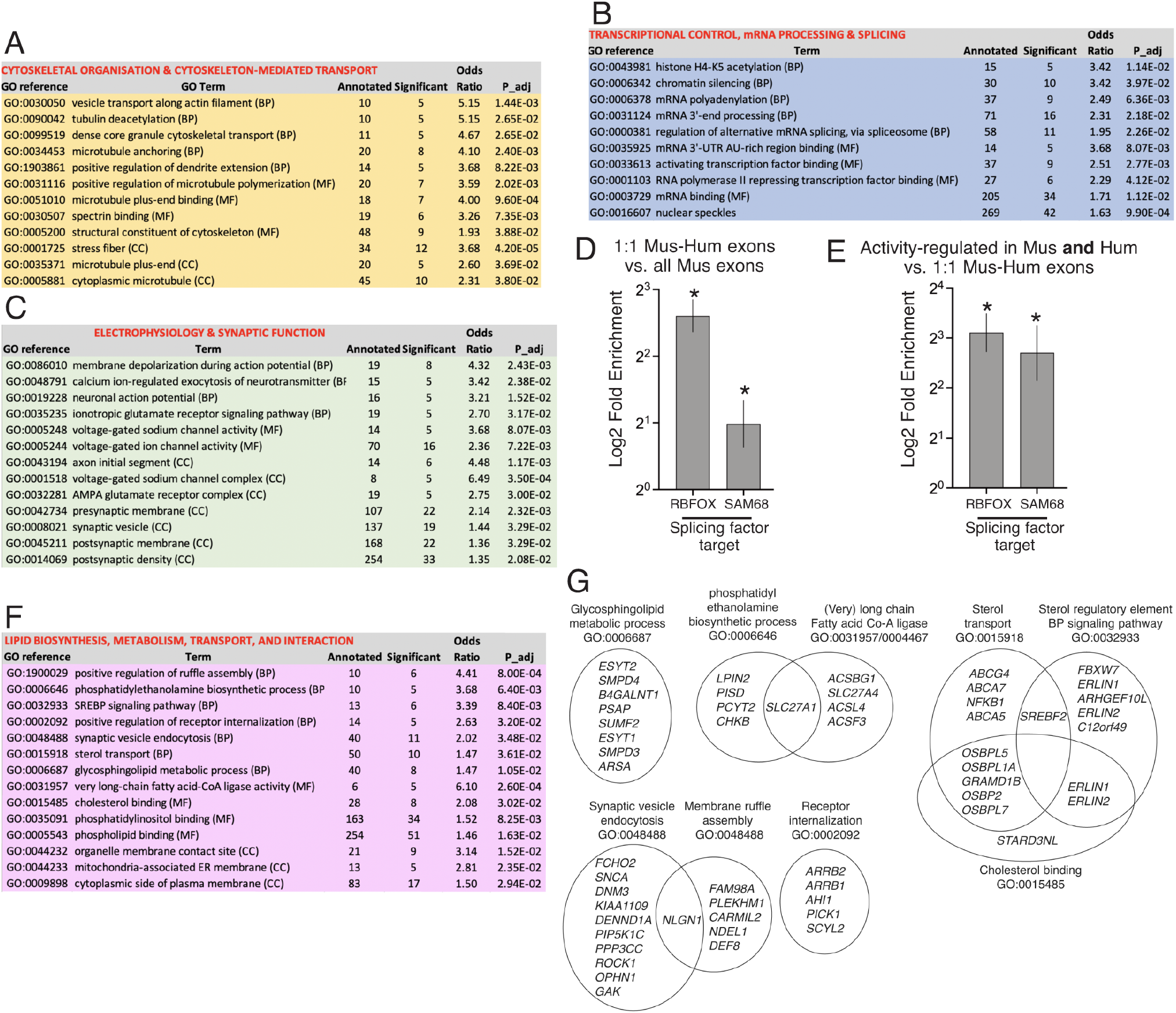
Ontology of genes subject to human-mouse conserved and human-specific activity-dependent exon usage. **A-C)** Selected GO terms are shown which are enriched (Fishers weighted p <0.05) in genes that have one or more 1:1 human-mouse orthologous exon which are subject to activity-dependent inclusion/exclusion in both Hum-ESC^CORT^ neurons and in one or more of our mouse cortical neuronal preparations (DIV4 and DIV10 Mus-PRIM^CORT^, mESC^CORT^-neurons). 782 genes contain exons that qualify as being regulated in a conserved manner by the above criteria, out of a background of 8039 genes (defined as 1:1 orthologous genes possessing ≥ 1 orthologous exons expressed in human and mouse neurons). The nature of the GO term is shown (BP: Biological Process; MF: Molecular Function; CC: Cellular Component). **D**,**E)** Enrichment tests were performed for RBFOX and SAM68 target cassette exon splicing events (Farini et al., 2020; Jacko et al., 2018) (see Methods). For (D) the presence of these target exons was compared between the set of all exons expressed in mouse neurons, and the set of exons expressed in mouse neurons for which there is a 1:1 human ortholog. For (E) the presence of these target exons was compared between the set of 1:1 orthologous exons subject to activity-dependent splicing in both human and mouse neurons (as per A-C), and the whole set of expressed 1:1 orthologous exons. **F**,**G)** Selected GO terms are shown which are enriched (Fishers weighted p <0.05) in genes that have a 1:1 human-mouse orthologue which are subject to activity-dependent alternative splicing in human neurons but not mouse neurons. In (G) the genes within selected GO terms that account for the enrichment are shown, and any genes in more than one GO term indicated by the overlapping nature of the Venn diagram. Note that while selected pathways are shown in this figure, all significantly enriched pathways are shown in Source_Data.xlsx.

We were also interested in identifying putative regulators of this conserved group of activity-dependent exons. Dynamic changes in exon usage in developing neurons are controlled by several RNA binding factors, prominent among them being the RBFOX proteins (1-3) and SAM68 (Farini et al., 2020; Furlanis and Scheiffele, 2018; Jacko et al., 2018), the latter particularly implicated in mediating activity-dependent exon usage (Farini et al., 2020; Iijima et al., 2011). Consistent with conserved importance of RBFOX and SAM68-regulated exons, their target exons (See Methods and (Farini et al., 2020; Jacko et al., 2018)) are enriched 6.1-fold (Rbfox) and 2.0-fold (Sam68) in orthologous exons compared to non-orthologous ones (Fig 3d). Moreover, even after allowing for this enrichment, orthologous exons exhibiting activity-dependent changes in FEI in both human and mouse neurons (subject to GO analysis in Fig. 3a-c) are further enriched in RBFOX targets by 8.6-fold and in SAM68 targets by 6.1-fold (Fig .3e). Thus, both RBFOX- and SAM68-regulated exons are both conserved in terms of gene structure and additionally form a significant element of conserved exons subject to activity-dependent exon inclusion in human and mouse neurons.

We also wanted to determine whether genes subject to human-specific activity-dependent exon inclusion were enriched in any specific biological processes. We studied genes that have a 1:1 human-mouse orthologue which are subject to human neuron-specific activity-dependent alternative splicing. “Human-specific” regulation was defined as genes that in Hum-ESC^CORT^ neurons had 1 or more exons whose fractional inclusion in the mature transcript changed >0.1 (up or down, p<0.05) upon KCl treatment, but whose mouse ortholog did not meet these criteria in any of the mouse neuronal preparations used in this study. 1104 genes were found to be subject to human-specific alternative splicing (by the criteria above) and were subject to ontological analysis. Biological processes and molecular function terms enriched in genes subject to human-specific activity-dependent alternative splicing are dominated by lipid biology (Fig. 3f,g). This includes gene sets involved in lipid/phospho-lipid biosynthesis, regulation of lipid biosynthesis (e.g. signaling to the sterol regulatory element) and lipid interaction, as well as lipid transport and processes sassociated with lipid bilayer dynamics such as synaptic vesicle endocytosis and membrane ruffle assembly. The prominence of a wide range of largely gene sets relating to lipid biology (Fig. 3f,g) is also clear when reviewing all pathways that are significantly enriched (see Source_Data.xlsx). The functional consequences of actvity-dependent AS of so many lipid pathway genes will require further investigation.

### Cis vs. trans determinants of basal and activity-dependent alternative splicing divergence

Having established that in forebrain neurons that both basal exon inclusion levels and activity-dependent changes in exon inclusion show a degree of evolutionary divergence between mouse and human, we wanted to determine whether this divergence was due to changes in cis-acting factors (i.e. DNA sequence) or trans-acting factors, such as splicing factors or signal transduction machinery. Neurons from the Tc1 transchromosomic mouse strain (O’Doherty et al., 2005), were used to assess the FEI of human chromosome 21 exons which have an orthologous mouse exon, whose FEI could be measured in the same RNA sample, employing our *in silico* species-specific RNA-seq read analysis (Qiu et al., 2018). The FEI of the human exons in Tc1 mouse neurons could be compared to the FEI of those same exons in a human cellular environment (i.e. hESC^CORT^-neurons). Of the 45 hCh21 exons which both had a 1:1 mouse orthologue and passed an expression level threshold, there was a significant correlation comparing FEI of human exons in Tc1 cortical neurons vs. Hum-ESC^CORT^ neurons (Fig. 4a), and (as expected) the corresponding mouse exons showed near-identical FEI in Tc1 cortical neurons vs. DIV10 Mus-PRIM^CORT^ neurons (Fig. 4b)

**Figure 4.**
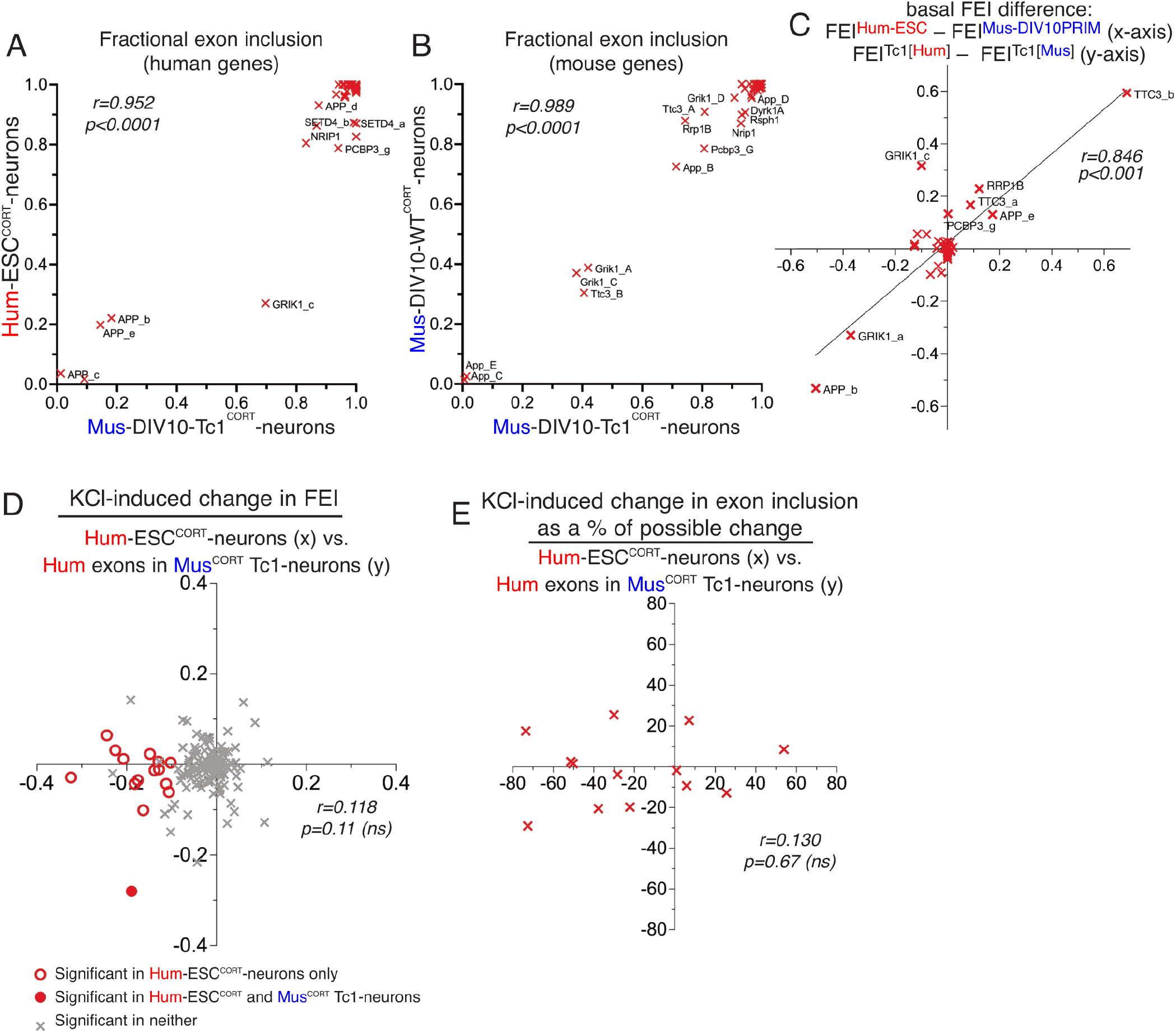
Cis vs. trans determinants of basal and activity-dependent alternative splicing divergence. **A)** FEI of hCh21 exons (with a 1:1 mouse/human ortholog) in Hum-ESC^CORT^ neurons vs. the FEI of the human exon in mouse Tc1 neurons. **B)** FEI of the mouse orthologs of the exons from Fig. 4a in mouse Tc1 neurons (x-axis) vs .FEI in DIV10 Mus-PRIM^CORT^ neurons. **C)** A comparison of the difference in basal FEI in orthologous exons within mouse Tc1 neurons compared to the difference in the same exons between mouse (DIV10 Mus-PRIM^CORT^) and human (hESC^CORT^) neurons. Correlation coefficient r is shown. D) A comparison of KCl-induced FEI changes in human Ch21 exons in Hum-ESC^CORT^ neurons vs. mouse Tc1 neurons. E) For alternatively spliced exons (0.8>basal FEI>0.2) the effect of KCl stimulation on FEI as a percentage of the maximum possible FEI change was compared between Hum-ESC^CORT^ neurons vs. mouse Tc1 neurons. For data points relating to this figure see Source_Data.xlsx.

We also wanted to determine whether human-mouse differences in FEI of orthologous exons observed when comparing the transcriptomes of human and mouse neurons were also observed when those exons were studied in the same cellular environment (mouse Tc1 neurons). We observed a good correlation between human-mouse differences in FEI when studying them in their own cellular environment vs. a common Tc1 cellular environment (Fig. 4c). This supports a model whereby FEI is dictated by cis-acting DNA elements and that evolutionary divergence of basal exon inclusion is driven at least in part by changes in cis-acting DNA elements.

We next investigated whether activity-dependent changes in FEI are also determined by cis-acting factors. Activity-dependent changes in FEI were assessed in human Ch21 exons within the mouse Tc1 neurons, and this was compared to changes of the same exons in Hum-ESC^CORT^ neurons. This revealed no significant correlation (Fig. 4c). We also performed additional comparisons of KCl-induced FEI differences calculated as a percentage of the maximum possible change restricting our analysis to orthologous exons where basal FEI was >0.2 and <0.8 as before (a similar approach to that taken in Fig. S2). This also revealed no correlation between activity-dependent changes in human exon inclusion in a human vs. mouse cellular environment (Fig .4d). There are 15 hCh21 exons (spanning 10 different genes) that undergo activity-dependent changes in inclusion in Hum-ESC^CORT^ neurons (FEI change ≥0.1 in any direction, p<0.05) and whose regulation could be studied in mouse Tc1 neurons. Only 1 out of those 15 human exons were controlled by KCl treatment of Tc1 neurons (Fig 4d). Thus, unlike basal exon inclusion levels which are relatively insensitive to cellular environment, activity-dependent changes in human gene exon inclusion appears to require a human cellular environment to be properly executed. These “trans-acting factors” may include the neuron’s activity-responsive signaling or splicing machinery. However, we cannot rule out that certain conserved alternatively spliced exons rely solely on cis-acting factors. Of the 15 activity-responsive human exons that could be analysed in Tc1 neurons none were putative RBFOX or SAM68 targets. It is possible that conserved exon targets of these factors are controlled by mouse splicing factors directed solely by cis-acting binding sites for these factors, although whether RBFOX, SAM68 and other factors (such as SRRM4) have cross-species activity is not clear.

### Concluding remarks

To conclude, our study indicates that there is significant conservation of both basal and activity-dependent exon usage between cortical-patterned human and mouse neurons. The classes of genes previously identified as being subject to activity-dependent alternative splicing in mouse neurons: synaptic, electrophysiology, cytoskeletal (Farini et al., 2020; Furlanis and Scheiffele, 2018; Iijima and Yoshimura, 2019; Jacko et al., 2018), also feature strongly in genes whose splicing is similarly regulated in human neurons. These genes whose activity-dependent splicing is conserved are also enriched in regulatory targets of splicing factors RBFOX and SAM68. However our study also supports the notion that there are both quantitative and qualitative differences in orthologous exon usage in human neurons, compared to their mouse ortholog. Moreover, differences in both basal exon usage and the activity-dependency of exon usage are apparent, although cis-acting sequences seem to be sufficient to drive the former, but not the latter. It is conceivable that the functional impact of neuronal activity on human forebrain neurons is different to those from mice, and that these differences may arise from alternative exon usage and not just differential regulation at the transcriptional level (Hardingham et al., 2018; Qiu et al., 2018). The prominence of lipid biology in genes with exons subject to human-specific control by neuronal activity is intriguing and provides a basis for further functional investigation.

## Acknowledgements

This work was funded by the UK Dementia Research Institute (GEH, SC) which receives its funding from DRI Ltd, funded by the UK Medical Research Council, Alzheimer’s Society, and Alzheimer’s Research UK. We acknowledge the work of all the authors of the studies who generated the primary data analysed in this study (Farini et al., 2020; Hasel et al., 2017; Jacko et al., 2018; Qiu et al., 2016; Quesnel-Vallieres et al., 2016).

## Data availability

RNA-seq data analysed is from the following our previously published work: E-MTAB-5489 (Qiu et al., 2016) and E-MTAB-5514 (Hasel et al., 2017) and published by others: GSE89984 (Quesnel-Vallieres et al., 2016). Code to enable species separation of mixed species RNA-seq data is available (http://biomedicalinformaticsgroup.github.io/Sargasso/) as described (Qiu et al., 2018). Numerical data used to generate the Figures are provided in file Source_Data_xlsx.

## Materials and Methods

### Splicing Analysis

We used RNA-seq data from the following accessions: E-MTAB-5489 (Qiu et al., 2016), E-MTAB-5514 (Hasel et al., 2017) and GSE89984 (Quesnel-Vallieres et al., 2016). Samples containing RNA-seq reads from only a single species were mapped to their respective genome using the STAR version 2.7.0f (Dobin et al., 2013); reads were mapped to the primary assemblies of the human (hg38) or mouse (mm10) reference genomes contained in Ensembl release 99 (Cunningham et al., 2022). Samples containing RNA-seq reads derived from both the human and mouse genomes (or single-species samples which were to be compared with these) were processed with Sargasso version 2.1 (Qiu et al., 2018) to disambiguate reads between the two species, using a conservative filtering strategy, to prioritise minimizing the number of read mis-assigned to the wrong species. In order to measure levels of exon inclusion, and differences in exon inclusion between experimental conditions, data were then processed with the differential splicing tool rMATS, version 4.1.0 (Shen et al., 2014), focusing on the “skipped exon” category of splicing events. Significance for differential inclusion events was generally defined as p < 0.05, average read count for exon inclusion or exclusion over all samples > 5, inclusion level difference > 0.1.

To match orthologous skipped exon events, the co-ordinates of mouse events were transformed from mm10 to hg38 co-ordinates using the command-line version of the UCSC liftOver tool (https://genome.ucsc.edu/cgi-bin/hgLiftOver). Mouse and human exon inclusion/exclusion events were then considered to be orthologous if the human co-ordinates and lifted mouse co-ordinates of the upstream exon end, downstream exon start, and the start and end of the alternatively spliced exon could be matched to within ten base-pairs.

### Enrichment analysis

Enrichment analyses were performed as follows. For gene ontology enrichments, a background gene set was constructed consisting of all human genes for which an event with average read count for exon inclusion or exclusion over all samples greater than 5 was tested for differential splicing, which had a 1:1 orthologous mouse gene, and for which the mouse gene had an event with average read count > 5 tested in at least one of the DIV4, DIV10 or mESC KCl vs basal comparisons. Then gene ontology enrichment was tested in (i) those human genes which had a significant differential splicing event (according to the definition above), for which the 1:1 orthologous mouse gene had a significant differential splicing event in at least one of the DIV4, DIV10 or mESC KCl vs basal comparisons; (ii) those human genes which had a significant differential splicing event for which the 1:1 orthologous mouse gene did not have a significant differential splicing event in any of the DIV4, DIV10 or mESC KCl vs basal comparisons. Gene ontology enrichment analyses were performed using topGO, R package version 2.42.0 (Shen et al., 2014).

At the level of splicing events themselves, enrichments for targets of particular splicing factors were tested using Fisher’s exact test. For each splicing factor three enrichment tests were performed: (i) in the background of all mouse events with average read count for exon inclusion or exclusion over all samples greater than 5 which were tested for differential splicing in at least one of the DIV4, DIV10 or mESC KCl vs basal comparisons, enrichment for those events with an orthologous human event with average read count greater than 5 which was tested for differential splicing; (ii) in the background corresponding to the enrichment set in (i), enrichment for those events which were significant in the human KCl vs basal comparison, and also in at least one of the mouse DIV4, DIV10 or mESC KCl vs basal comparisons; (iii) in the background corresponding to the enrichment set in (i), enrichment for those events which were significant in the human KCl vs basal comparison, but not in any of the mouse DIV4, DIV10 or mESC KCl vs basal comparisons (these latter two event-level enrichment tests correspond to the two gene-level enrichment analyses above). Enrichment tests were performed for (i) Rbfox target cassette exon splicing events determined by RNA-seq profiling after 10 days of maturation in Rbfox triple KO vs WT neurons as reported (Jacko et al., 2018); (ii) the union of Sam68 cassette exon splicing events determined by RNA-seq from Sam68 KO vs WT neurons at P1 and P10 (Farini et al., 2020).

---

**Figure S1.**
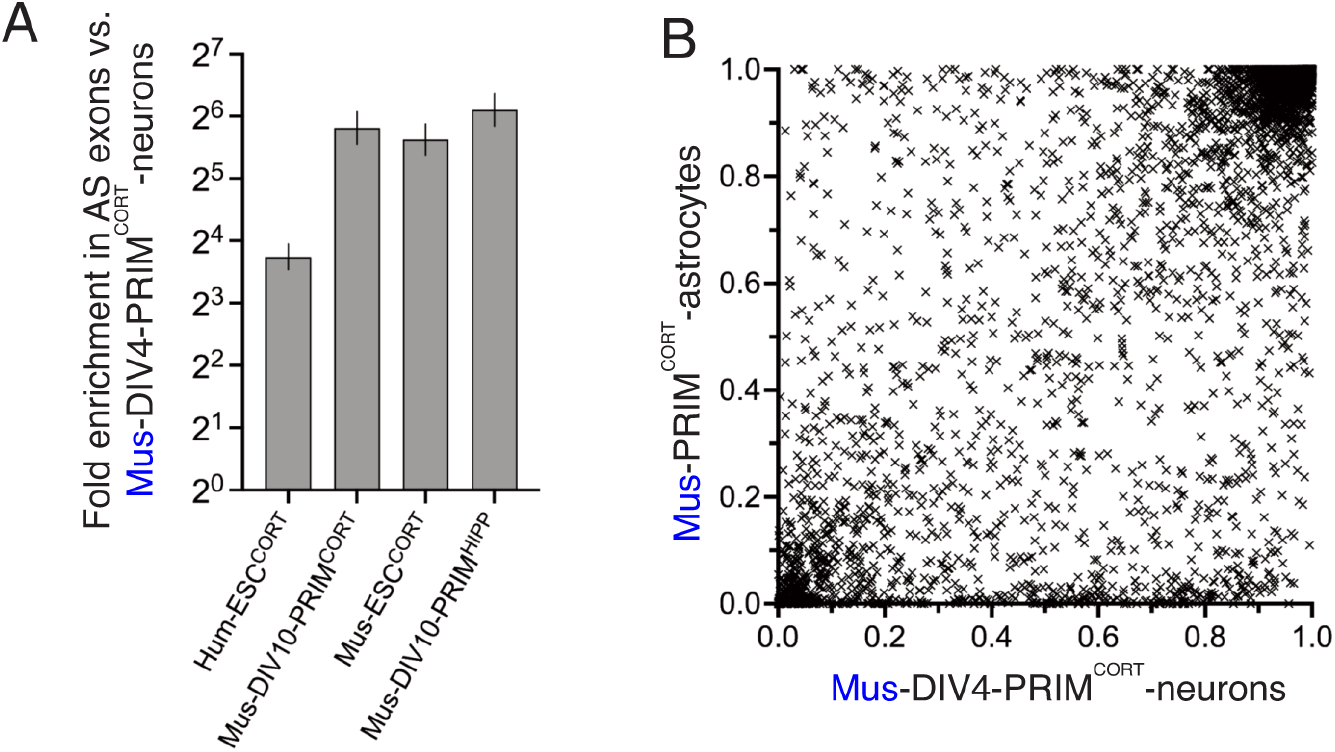
**A)** Fold enrichment of exons classed as alternatively spliced in DIV4 Mus-PRIM^CORT^ neurons in exons which are also classed as alternatively spliced in the indicated neuronal cell types (defined as (0.8>mean FEI>0.2, n=3). Error bars indicate the 95% confidence limits of the enrichment factor. In all cases p<0.0001 (Fisher’s exact test). **B)** Fractional exon inclusion (FEI) in DIV4 Mus-PRIM^CORT^ neurons plotted against the corresponding FEI in mouse cortical astrocytes (mean FEI, n=3) for all exons analysed in Fig. 1. For data points relating to this figure see Source_Data.xlsx.

**Figure S2.**
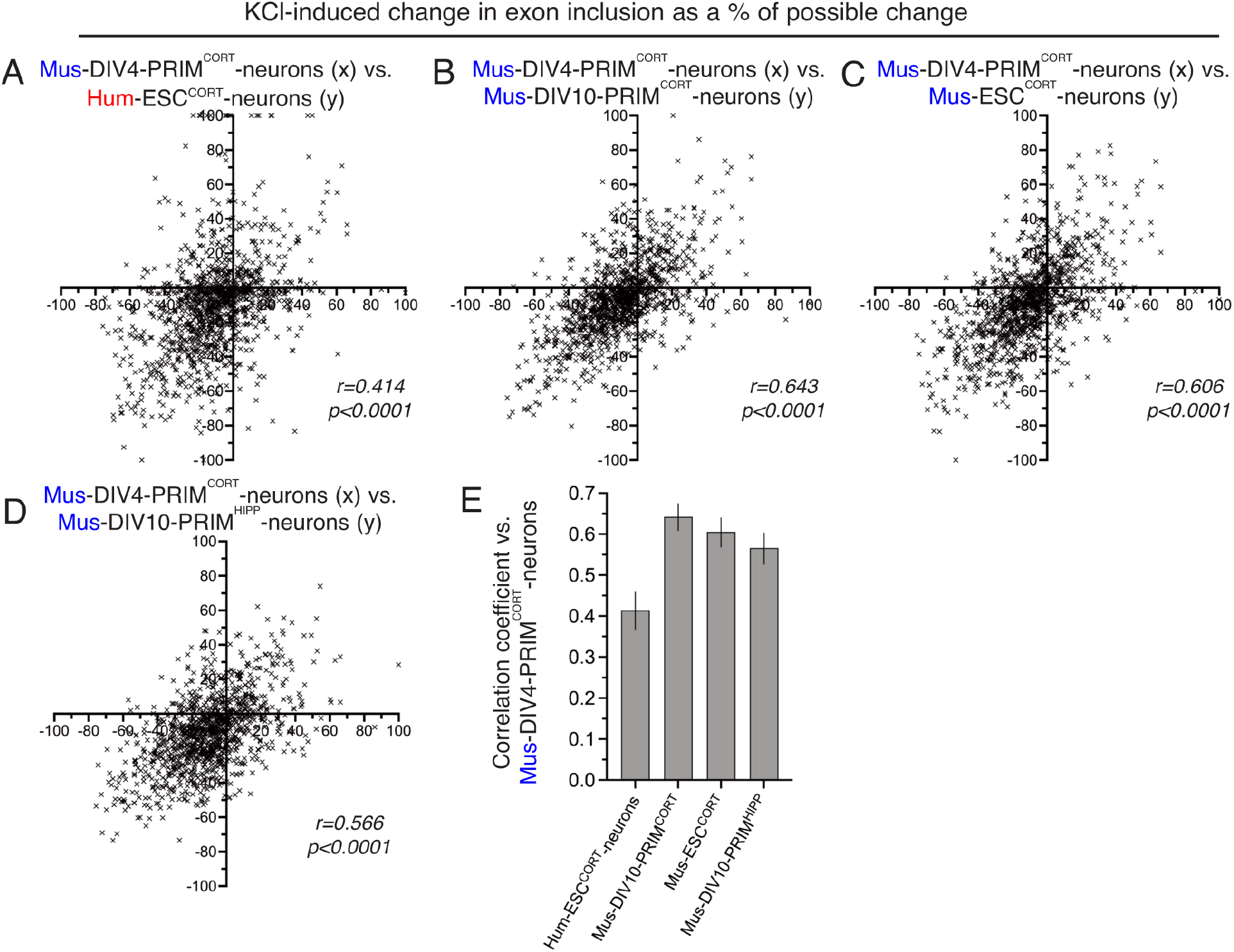
**A-D)** For exons classed as alternatively spliced (0.8>FEI>0.2) in DIV4 Mus-PRIM^CORT^ neurons, the effect of KCl stimulation on FEI was calculated as a percentage of the maximum possible FEI change and plotted (x-axis) against the corresponding value for the other cell types (y-axis) as indicated. **E)** Correlation coefficients for the comparisons made in A-D.. Error bars indicate the 95% confidence limits and in all cases p<0.0001. For data points relating to this figure see Source_Data.xlsx.

**Figure S3.**
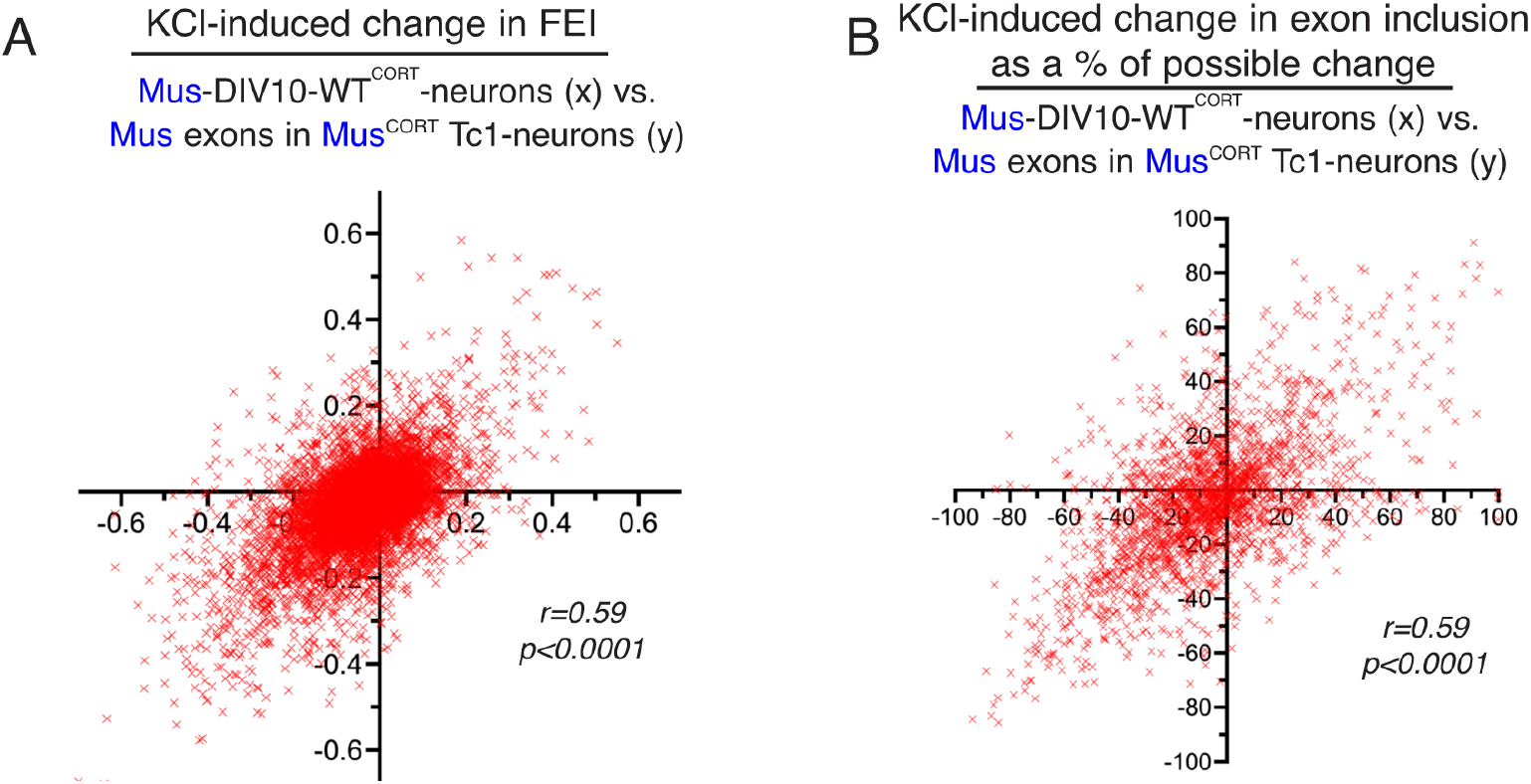
**A,B)** A comparison of KCl-induced changes in mouse exon FEI in Tc1 mouse neurons vs. wild-type mouse neurons. Both absolute FEI change is compared (A) as well as a percentage of the maximum possible FEI change (B) for alternatively spliced exons (0.8>basal FEI>0.2). For data points relating to this figure see Source_Data.xlsx.

